# Hyaluronidase unlocks sequestered CEACAM5 and improves CAR T-cell therapy in colorectal cancer

**DOI:** 10.1101/2024.08.29.610222

**Authors:** Debasis Banik, Christopher Ward, Ziwei Zhang, Soura Chakraborty, Daniel Heraghty, Prasanna Suresh, Bing Li, Shekhar Kedia, Simon J. Davis, James P. Roy, Michael A. Chapman, Bidesh Mahata, David Klenerman

## Abstract

Chimeric antigen receptor (CAR) T-cell therapy has shown unprecedented success in haematological cancers but faces challenges in solid tumours. Although carcinoembryonic antigen-related cell adhesion molecule 5 (CEACAM5) is differentially expressed in many solid tumours, anti-CEACAM5 CAR T-cells are ineffective. Here, we have studied the interaction of CEACAM5 targeting primary CAR T-cells with colorectal cancer (CRC) cells using fluorescence microscopy. We found that CRC cell’s glycocalyx is much deeper than that of the CAR T cell causing delayed activation. Oscillating calcium fluxes, indicative of non-sustained CAR T cell activation and reduced cytotoxicity, were observed when CAR T cells interacted with CRC cells, which increased with increasing cell-seeding time. Imaging revealed that this effect correlated with a progressive loss of accessible CEACAM5 antigen on the CRC cell surface, possibly due to their sequestration in the intercellular junction, rendering CAR T cell engagement less effective. Local proteolytic treatment with trypsin to disrupt the CRC cell monolayer, using a micropipette, increased CEACAM5 availability, decreased glycocalyx thickness, and restored sustained CAR T cell calcium fluxes, increasing the killing of CRC cells. Similar enhanced interaction was observed after treatment of CRC cell monolayer with hyaluronidase, approved for use in humans. We observed limited availability of CEACAM5 on human colorectal cancer tissues, whereas treatment with trypsin or hyaluronidase increased accessibility. Our results reveal why CAR T cells targeting CEACAM5 are ineffective and suggest possible routes to improved therapy for CRC.

## INTRODUCTION

T cells are an essential part of the adaptive immune system. Each T cell expresses T-cell receptors (TCRs) which recognise peptide major histocompatibility complexes (pMHCs) on the surface of antigen presenting cells (APCs). In contrast, synthetically designed chimeric antigen receptors (CARs) originally comprised target-binding single chain variable fragments (scFvs) of antibodies connected to an intracellular signaling domain, i.e., that of the CD3ζ subunit of the TCR, via a transmembrane region. Additional signaling domains, i.e., from CD28 and 4-1BB contribute to later generations of CARs. Recently, CAR T-cell therapy has been approved by the United States Food and Drug Administration (FDA) for the treatment of B cell acute lymphoblastic leukemia (ALL), with several products now in use in the clinic (*1–4*). However, the same therapy is ineffective in solid tumours, including colorectal cancer (CRC) (*5–8*). CRC, the third most common global cancer, is responsible for ∼10 % of total cancer incidences and deaths in 2020, with ∼ 30 months of median overall survival for metastatic CRC (mCRC) patients (*9*).

Understanding CAR T cell activity and limitations is essential for generating better therapies. CAR T cells require higher antigen concentrations than native T cells for activation (*10*). Lck independent CAR T cell activation was shown to be driven by the CD28 signaling domain and the SRC family kinase Fyn (*11, 12*), leading to the formation of a non-classical and potent immunological synapse generating rapid cytotoxicity (*13*). Recently, Xiao et al. showed that the exclusion of the large phosphatase CD45 at the anti-CD19 CAR T-cell/Raji B cell interface, used as a model system, is intrinsic to CAR T cell activation, likely due to the phosphatase exclusion favouring net CAR phosphorylation by kinases, as proposed by the ‘kinetic-segregation’ (KS) model (*14, 15*).

Carcinoembryonic antigen-related cell adhesion molecule 5 (CEACAM5) is overexpressed in many cancer types including lung, breast (luminal A & B), pancreatic tumours, and CRC. There has been little progress treating these tumours with CAR T cells for multiple reasons, such as poor accessibility of the cancer cells to the CAR T cells, poor persistence of the CAR T cells, and respiratory toxicity (*16*). Here, we investigate the interaction of anti-CEACAM5 CAR T-cells with CEACAM5^+^ CRC cell lines in vitro and with advanced model bilayers, using imaging methods, established previously (*17*). We observed that the unavailability of CEACAM5 on CRC cell monolayers and the deep glycocalyx of CRC cell play crucial roles in reducing the effectiveness of CAR T-cell/CRC cell interactions. We also observed enhanced interaction and killing of CRC cells when we locally treated the CRC cell monolayers with a proteolytic enzyme, trypsin or glycocalyx-degrading enzyme, hyaluronidase. CEACAM5 was also concealed in human colorectal cancer tissues, but increased availability was observed following treatment with either trypsin or hyaluronidase. Our results suggest routes to improved immunotherapy of CRC with hyaluronidase/CAR T-cells.

## RESULTS

### Impaired activation of anti-CEACAM5 CAR T-cell in colorectal cancer models

Jurkat and primary human T-cell derived anti-CEACAM5 CAR T-cells were prepared from clone hMN-14 which binds to the A3B3 domain of CEACAM5 (Fig. S1A-F) (*18, 19*). CEACAM5^-^ (RKO as control) and CEACAM5^+^ (LS-174T & LS-1034) CRC cells were used in our experiment (Fig. S1G-I). To establish baseline CAR T-cell functionality, we first validated anti-CEACAM5 Jurkat CAR T-cells on our “second generation” supported lipid-bilayer system (SLB2), which presents ligands for the small and large adhesion proteins CD2 (i.e., CD58) and LFA-1 (i.e., ICAM-1), alongside the major antigen presenting cell (APC) glycocalyx elements, CD43 and CD45 (∼ 40 nm in length), human CEACAM5/CD66e protein as an antigen for the CAR T-cell, and gp100-MHC, an irrelevant peptide-MHC used to block free nickellated lipids.

We prepared CEACAM5 positive (CEACAM5^+^) and CEACAM5 negative (CEACAM5^-^) SLB2s. CAR T-cells labelled with Fluo-4-AM (a calcium indicator) and CellMask Deep Red (a plasma membrane stain) were then allowed to interact with the SLB2s. We observed that CAR T-cells formed close contacts by excluding CD45/CD43 on the bilayer and produced robust calcium fluxes on the CEACAM5^+^ SLB2s (Movie S1). On CEACAM5^-^ (control) SLB2s, transient interactions and no calcium fluxes were observed (Fig. S2A & Movie S1). This result shows that CAR T-cells are highly sensitive to the presence of human CEACAM5 protein. Interestingly, we observed ∼ 40 % average exclusion of CD45/CD43 by CAR T-cells (Fig. S2B), similar to wild-type Jurkat and primary T-cells in our previous study (*17*), suggesting similar interactions with the SLB2 system.

Fig. 1 shows early-stage imaging of the anti-CEACAM5 CAR T-cell/CRC cell interaction using brightfield and epifluorescence calcium imaging. We sought to mimic tumour growth-related changes by plating out CRC cells for 24 and 48 hrs. Fig. 1A shows the experimental scheme for the cell-cell assay wherein Fluo-4-AM pre-labelled CAR T-cells interact with the LS-174T monolayer and produce calcium fluxes. Noting the off-target respiratory distress experienced by colon cancer patients within 15 minutes of CAR T-cell infusion (*20*), we decided to focus on the first one hour of CAR T-cell/LS-174T interaction using simultaneous measurement of calcium fluxes and brightfield images.

**Fig. 1:**
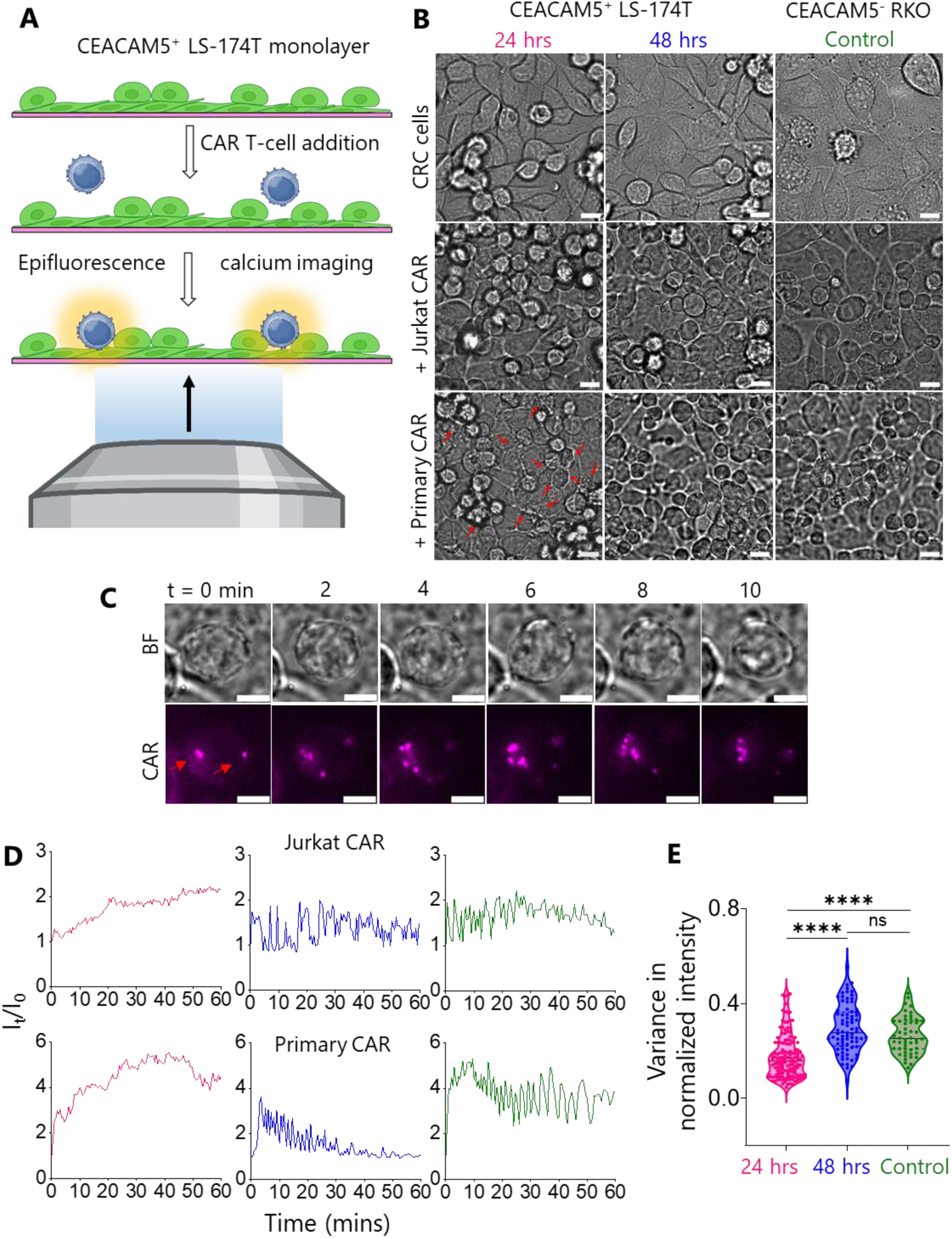
CAR T-cell/CRC cell interaction. (A) Cartoon shows CAR T-cells interacting with CEACAM5^+^ LS-174T cell monolayer. Live T-cell/tumour interaction was monitored using epifluorescence calcium and brightfield imaging. (B) Brightfield images show differently seeded LS-174T (pink, blue: 24 and 48 hrs, respectively) and CEACAM5^-^ RKO monolayers (olive: 24 hrs) interacting with Jurkat and primary CAR T-cells. Red arrows indicate killing of LS-174T cells by primary CAR T-cells. Scale bars are 10 μm. The same colour code will be used hereafter throughout all figures. (C) Time-series showing CAR accumulation (red arrows) while Jurkat CAR T-cells interacting with 24 hrs seeded LS-174T monolayer. Scale bars are 5 μm. (D) Representative epifluorescence calcium flux colour coded traces of Jurkat and primary CAR T-cells while interacting with LS-174T and RKO monolayers. (E) Violin plot shows variance in normalized calcium flux intensity of CAR T-cells on CRC cell monolayers. Data shown in D & E were pooled from 6-7 and 3-4 independent experiments of Jurkat CAR T-cell/CRC cell and primary CAR T-cell/CRC cell interactions, respectively with n = 145 (24 hrs), 77 (48 hrs), and 50 (Control). The black lines indicate the median which were compared using two-sided Student’s t-test. ****P <0.0001, ns not significant.

Firstly, we monitored the interaction using a 20X objective lens and found that the Jurkat CAR T-cells produced sustained calcium fluxes (red arrows in Movie S2) with early-seeded LS-174T monolayer (24 hrs), whereas, no sustained flux was observed following longer seeding of the monolayers (48 hrs). The triggering of CAR T-cells was substantially delayed on early-seeded LS-174T monolayers compared to the SLB2 (peak activation longer than one hr vs. within 3 mins. in Fig. S2C). We repeated the same experiment with LS-1034, a second CEACAM5^+^ CRC cell line, and found almost no T-cell triggering, even for the early seeding condition (Movie S3).

To investigate signaling on LS-174T monolayers in more details, we used a 60X objective lens. Brightfield images of LS-174T and the RKO monolayer (24 hrs) interacting with Jurkat CAR T cells are shown in Fig. 1B. CAR accumulation was observed as bright puncta formation during interaction with early-seeded LS-174T monolayer (Fig. 1C). Representative calcium flux traces are shown in Fig. 1D and Movies S4, S5. The increased and sustained calcium flux changed to oscillatory fluxes with increasing seeding time. Interestingly, similar oscillatory calcium fluxes were also observed on the RKO monolayer, suggesting that at the later seeding time-points there is very little antigen recognition. Fig. 1B (3^rd^ row, red arrows) and Movie S6 reveal extensive killing of the early-seeded LS-174T cells by primary CAR T-cells. In contrast, within the first one hour of monitoring, no killing was observed for longer-seeded LS-174T and control monolayers. We present more calcium flux data Movies S7, S8 to support our observations. We determined the variance of calcium flux fluctuations (Fig. 1E). The variance in the normalised calcium flux fluctuations was substantially lower for T-cells interacting with the early-versus the longer-seeded LS-174T and control monolayers. This result confirmed that significant increases in calcium flux oscillation accompany the early- to longer-seeded LS-174T transition, with the latter condition being very similar to the control monolayer.

In summary, whereas anti-CEACAM5 CAR T-cells responded to early-seeded CRC cell monolayers, decreased activation was observed with increasing seeding time. Therefore, consistent with clinical observations (*16*), our results show that anti-CEACAM5 CAR T-cells respond poorly to CRC.

### The deepened glycocalyx of colorectal cancer cells creates a significant physical barrier to CAR T cell engagement

The glycocalyx is a critical surface coating on the surface of all cells that modulates cell-cell interactions and can influence immune surveillance. The glycocalyx comprises dense arrays of negatively charged, heavily glycosylated proteins and polysaccharides, that forms a physical barrier to cell contact (*21*). In cancer cells the glycocalyx increases in depth, making it more difficult for T-cells to kill cancerous cells, especially solid tumours (*22, 23*). Therefore, it was of interest to test whether the decreased CAR T-cell/LS-174T interaction is associated with the presence of a deep glycocalyx on the tumour cell.

To measure the glycocalyx, CAR T-cells and early- and longer-seeded LS-174T cells were fixed and labelled with HMSiR-conjugated wheat germ agglutin (WGA-HMSiR). WGA binds to the N-acetylglucosamine residues of the glycocalyx. We measured the glycocalyx depth using resPAINT super-resolution (SR) imaging (*24*). Fig. 2A-C show 2D SR images of T-cell and LS-174T glycocalyces, which are ∼ 100 nm and ∼ 400 nm in depth, respectively. Fig. 2D confirms that the LS-174T cell glycocalyx is significantly deeper than that of the CAR T-cells.

**Fig. 2:**
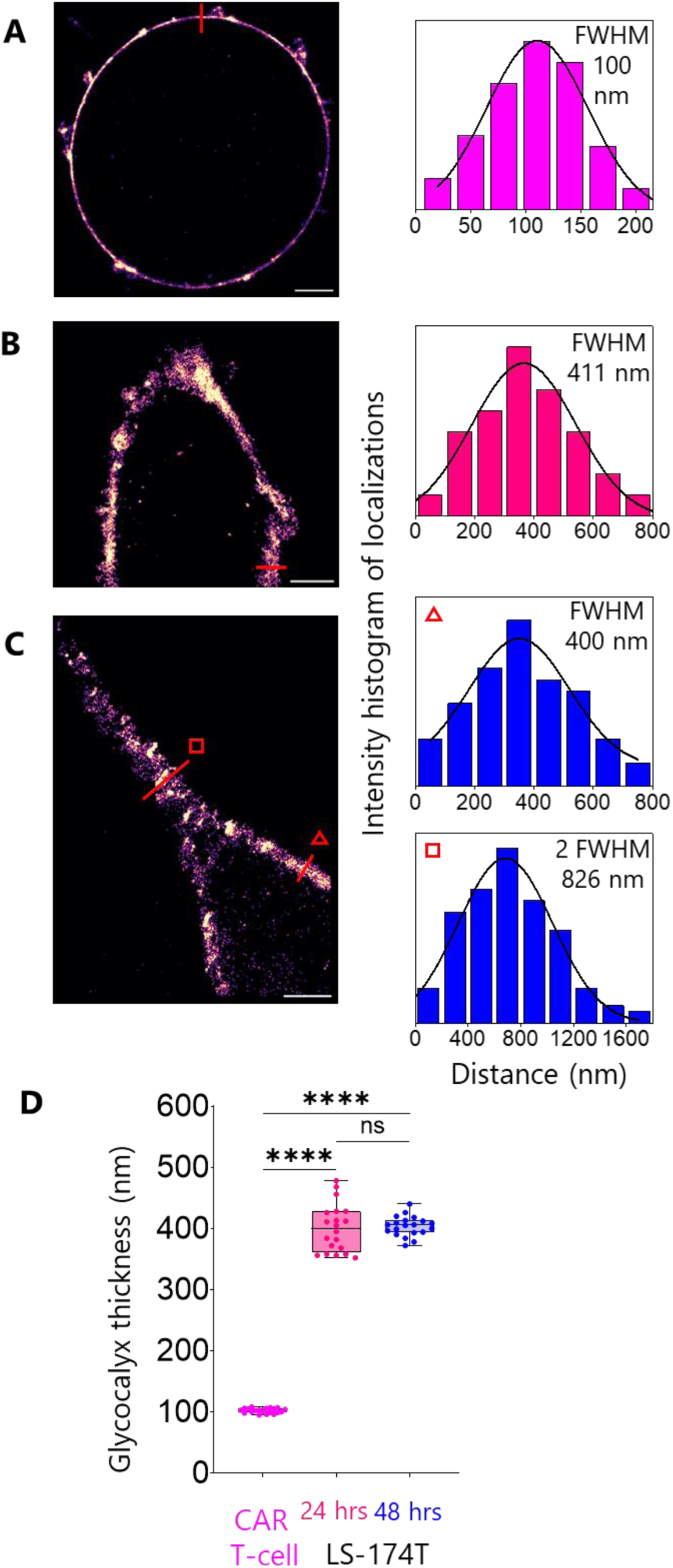
Super-resolution (resPAINT) images of the glycocalyx. Representative 2D images of (A) CAR T-cell and (B-C) LS-174T cell (24 and 48 hrs seeded, respectively) glycocalyces labelled with WGA-HMSiR. Scale bars are 2 µm. Colour coded line plots of 400 nm width as indicated in A-C are shown in front of images. Colour code for T-cell is magenta. Glycocalyx thickness is shown at two locations of C (triangle and square indicate 1*glycocalyx and 2*glycocalyx, respectively). Glycocalyx thickness was determined as the FWHM of the Gaussian fit. (D) Boxplot shows the glycocalyx thickness of CAR T-cell vs. LS-174T cell. In each case, data points (n = 20) were extracted by measuring thickness at multiple locations of three individual images collected from three independent experiments. Medians were compared using two-sided Student’s t-test. ****P < 0.0001, ns not significant.

Interestingly, no significant variation in glycocalyx depth was observed between early- and longer-seeded LS-174T cells (Fig. 2D), indicating that the differential CAR T cell reactivity observed between these conditions cannot be attributed to changes in glycocalyx dimensions alone. This suggests that while the substantial glycocalyx disparity represents a physical barrier to effective CAR T cell engagement with tumour cells, additional factors likely contribute to the seeding-dependent differences in CAR T cell activation and cytotoxicity.

These findings establish that the glycocalyx of CRC cells creates a formidable spatial barrier that must be overcome for effective immunotherapeutic targeting, providing insight into possible mechanism underlying the limited efficacy of CAR T cell therapy against solid tumours.

### CEACAM5 sequestration at tumour cell-cell junctions mediates immune evasion from CAR T cell recognition

To explore the reasons for the decrease in CAR T-cell/LS-174T reactivity with increasing seeding time, we performed antigen mapping on the LS-174T monolayer using fluorescently labelled anti-CEACAM5 antibody. Strikingly, CEACAM5 distribution on LS-174T cells exhibited a temporal pattern of sequestration. In minimally established cultures (4 hrs post-seeding), CEACAM5 was abundantly available across the entire cell surface (Fig. 3A, top row). On early-seeded monolayer, CEACAM5 redistributed predominantly to intercellular junctions with heterogeneous expression (Fig. 3A, second row), a pattern commonly observed for solid tumour antigens. Notably, in longer-seeded monolayers, CEACAM5 became increasingly inaccessible, with staining patterns resembling the CEACAM5-negative RKO controls (Fig. 3A, third and fourth rows). Quantitative analysis confirmed significant reduction in CEACAM5 accessibility on transitioning from early-to late-seeded LS-174T monolayers and the latter one showed no significant difference to RKO (Fig. 3B).

**Fig. 3:**
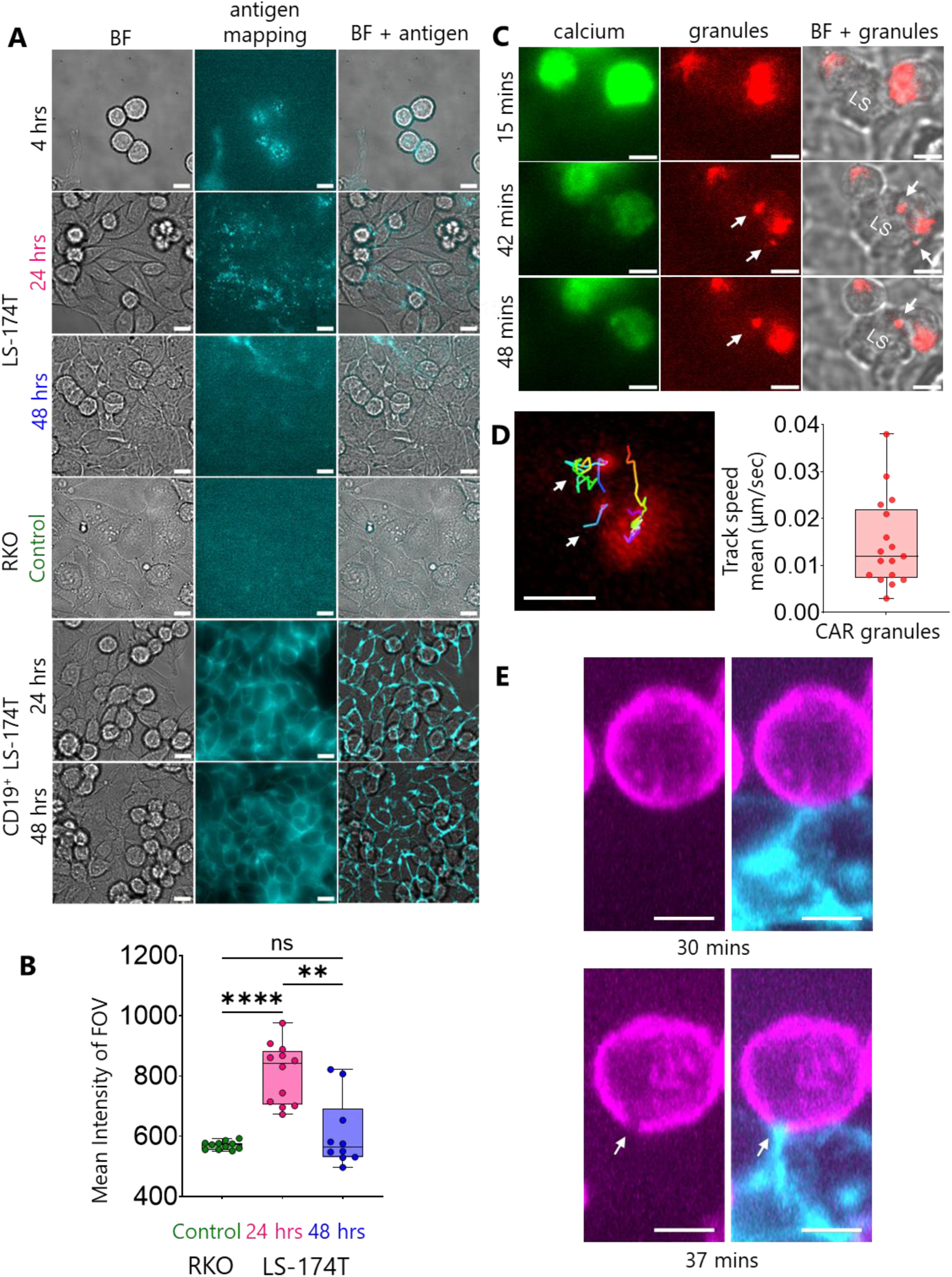
Antigen mapping, granules recruitment and CD45 exclusion while CAR T-cell/CRC cell interaction. (A) Middle column shows CEACAM5 and CD19 mapping on LS-174T, RKO and CD19^+^ LS-174T cells/monolayers, respectively at different seeded conditions mentioned on the left. The left and right columns show brightfield (BF) and merged (brightfield and antigen) images, respectively. For each seeding condition, images are representative of n = 10-15 FOVs. (B) Quantification of mean CEACAM5 intensity of FOV for 24 hrs (n = 12), 48 hrs seeded (n = 10) LS-174T and RKO (n = 11) monolayers. Medians were compared using two-sided Student’s t-test. ****P < 0.0001, **P = 0.0054, ns not significant. (C) Time series of calcium flux (left column) and granules recruitment (white arrows in middle and right columns) images show the sequence of events during primary CAR T-cell/LS174T cell (24 hrs seeded) interaction. Results are representative of n = 9 events collected from 3 individual FOVs. (D) Tracks (white arrows) and track speed mean (boxplot) of CAR granules. (E) Jurkat CAR T-cell (magenta)/LS-174T cell (cyan) conjugates at 30 and 37 mins after mixing. CD45 exclusion (white arrows) at cell/cell interface was captured using epifluorescence selective plane illumination microscopy (eSPIM) and representative of n = 12 FOVs. Scale bars are 10 μm and 5 μm in A, C, D and E, respectively.

This progressive antigen sequestration correlated with functional impairment, as evidenced by significantly increased T cell motility and track straightness on longer-seeded LS-174T monolayers compared to early-seeded cultures (Fig. S3A-D & Movie S9). To determine whether this sequestration phenotype was antigen-specific or a general characteristic of CRC cells, we examined the distribution of two additional clinically relevant antigens. We engineered CD19-expressing LS-174T cells (Fig. S3E-F) and examined endogenous HER2 expression. In contrast to CEACAM5, both CD19 and HER2 maintained homogeneous cell-surface distribution regardless of seeding duration (Fig. 3A, bottom rows; Fig. S3H-I).

Furthermore, anti-CD19 CAR T-cells exhibited robust calcium signalling against longer-seeded CD19⁺ LS-174T monolayers (Fig. S3G), confirming that CEACAM5 sequestration represents a distinctive immune evasion mechanism rather than a general feature of antigen presentation in maturing CRC cultures.

To characterize the functional consequences of CEACAM5 accessibility, we monitored primary CAR T cell interactions with early-seeded LS-174T monolayers using simultaneous calcium and lysosomal imaging. By labeling T-cells with Fluo-4-AM and lysotracker red, we observed calcium fluxes (∼ 15 mins after CAR T addition) preceding the recruitment of granules (∼ 40-50 mins) at the immunological synapse formed by the CAR T-cells with LS-174T cells (Fig. 3C & Movie S10). We analysed the recruitment trajectories of the granules (Fig. 3D & Movie S11) and found the mean speed of recruitment varies between 0.005 to 0.04 μm/sec. Davenport et al. did similar analyses of primary CAR T-cells targeting HER2 and found granule recruitment speeds of ∼ 0.10 μm/sec (*13*). This implies that granule recruitment speed is target-antigen dependent.

After observing CD45/CD43 exclusion on the SLB2 (Movie S1), we sought to image the same phenomenon at the CAR T-cell/tumour cell interface. Early-seeded LS-174T monolayers and Jurkat CAR T-cells were labelled separately with anti-epithelial cell adhesion molecule (EpCAM) antibody-Alexa 647 and Gap8.3-Alexa 488 (anti-CD45 antibody) conjugates, respectively, and then allowed to interact for 30 mins. Fig. 3E shows late stage CD45 exclusion on T-cells (white arrows) at the CAR T-cell/LS-174T interface using a home-built epifluorescence selective plane illumination microscope (eSPIM).

### Enzymatic remodelling of the tumour cell intercellular junctions increases CEACAM5 accessibility and enhances CAR T cell function

The ineffectiveness of CEACAM5-targeting CAR T cells against CRC may be attributed to two key barriers we identified: antigen sequestration at intercellular junctions (Fig. 3A, B & Fig. S4A, S4B) and the deep glycocalyx of CRC cells (Fig. 2). To address these limitations, we conceived the idea of treating longer-seeded LS-174T monolayers with the dissociating enzyme trypsin to re-expose the CEACAM5 and reduce the physical barrier of the glycocalyx.

To address CEACAM5 sequestration at intercellular junctions, we developed a localised trypsin delivery system using a micropipette (Fig. 4A). This approach selectively disrupted cell-cell adhesions in the upper layer of longer-seeded LS-174T monolayers without compromising monolayer integrity (Movie S12). Calcium flux analysis revealed sustained signaling in anti-CEACAM5 CAR T cells interacting with trypsin-treated LS-174T monolayers, mirroring responses observed in early-seeded cultures (Fig. 4B & Movie S13, S14). Control experiments using CEACAM5-negative RKO monolayers showed unchanged oscillatory calcium fluxes (Fig. 4B), confirming treatment specificity. Variance analysis demonstrated significant reduction in calcium flux oscillations after trypsin treatment, comparable to early-seeded conditions (Fig. 4C).

**Fig. 4:**
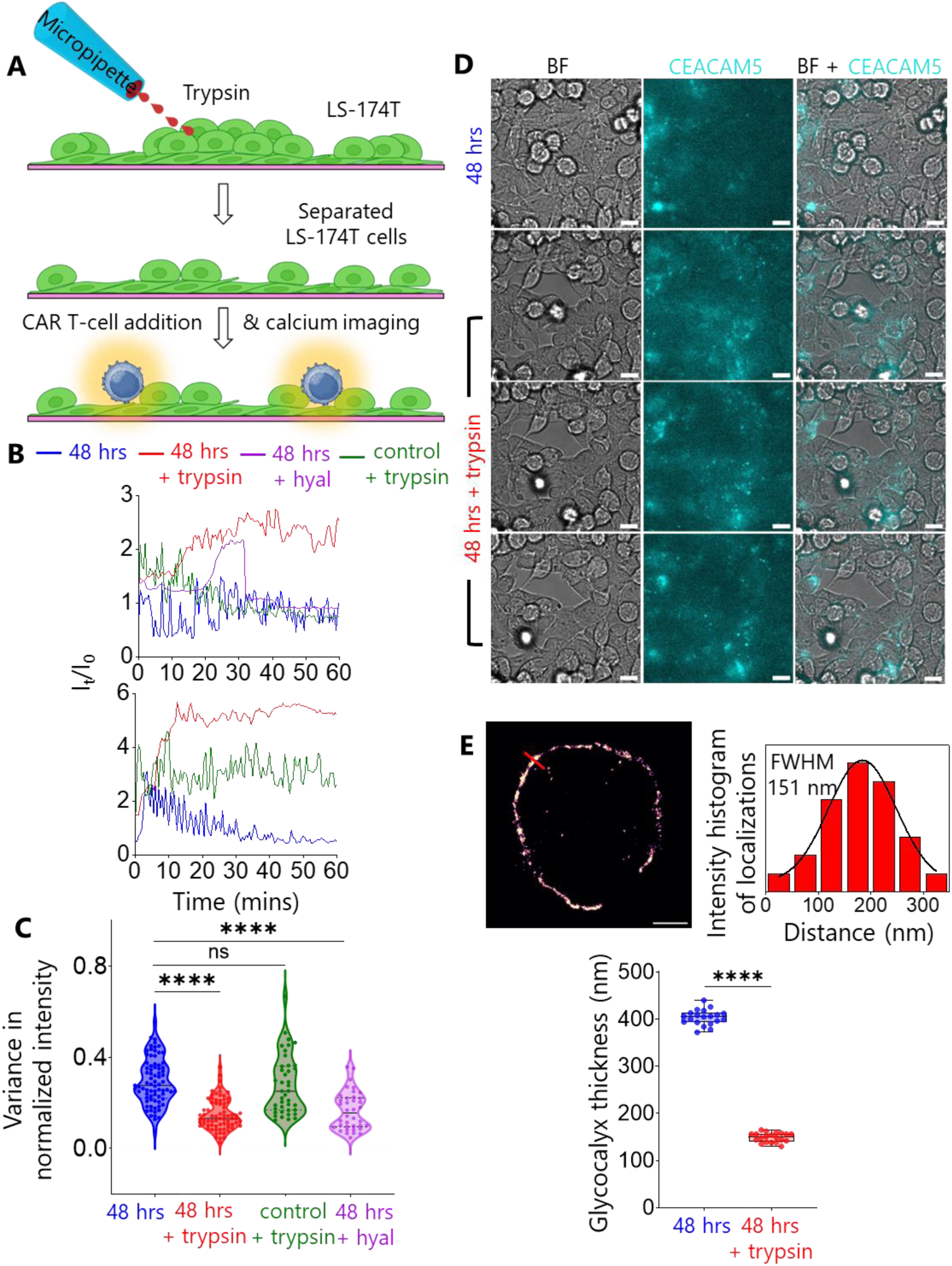
Trypsin and hyaluronidase (hyal) treatment to improve CAR T-cell/CRC cell interaction. (A) Cartoon shows trypsin treatment on 48 hrs seeded LS-174T monolayer using a micropipette to dissociate cells followed by CAR T-cell addition and epifluorescence calcium imaging. (B) Calcium flux traces of CAR T-cells before (48 hrs), after trypsin (48 hrs + trypsin, red) and hyaluronidase (48 hrs + hyal, purple) treatment on LS-174T monolayer, and after trypsin treatment on RKO (olive) monolayers. The y-axis of the graphs is shifted from 1 for better comparison. Top and bottom graphs indicate Jurkat CAR and Primary CAR, respectively. (C) Violin plot shows comparison of variance in normalized calcium flux intensity of CAR T-cells on CRC cell monolayers before and after trypsin, hyaluronidase treatments with n = 77 (48 hrs), 74 (48 hrs + trypsin on LS-174T), 41 (48 hrs + trypsin on RKO) and n = 28 (48 hrs + 25 U/ml hyaluronidase on LS-174T). (D) CEACAM5 mapping (middle row) on LS-174T monolayer before and after trypsin treatment. Scale bars are 10 μm. (E) 2D resPAINT image of LS-174T cell glycocalyx in presence of trypsin and corresponding line plot. Scale bar is 2 μm. Boxplot shows the comparison of glycocalyx thickness of LS-174T cell before and after trypsin treatment with n = 20 in each condition. The medians in C and E were compared using two-sided Student’s t-test. ****P < 0.0001, ns not significant.

Immunofluorescence mapping revealed rapid CEACAM5 re-availability at the intercellular junction within 20 minutes post-trypsin application, to the extent observed for the early-seeded condition (Fig. 4D). Super-resolution resPAINT imaging showed significant reduction of glycocalyx thickness to ∼ 150 nm in the presence of trypsin, while creating discontinuous glycocalyx patches (Fig. 4E).

We tested trypsin treatment on early-seeded LS-174T monolayers and observed faster triggering of Jurkat CAR T-cells measured as calcium fluxes, versus the untreated condition (Fig. S2C). CEACAM5 mapping showed increased availability after treatment (Fig. S4C).

This dual mechanism, antigen re-availability and glycocalyx thinning, explained restored CAR T cell activation kinetics, with primary CAR T-cells achieving target cell killing within one hr on treated monolayers versus no cytotoxicity in controls (Movie S15).

### Hyaluronidase pretreatment overcomes glycocalyx-mediated resistance to anti-CEACAM5 CAR-T cells

We investigated whether the improved CAR T-cell/LS-174T interaction is also observed when we apply glycocalyx specific enzyme to reduce/remove the depth or density of the glycocalyx of LS-174T cells. Hyaluronidase degrades hyaluronic acid, a component of the glycocalyx. We treated longer-seeded LS-174T monolayers with 25 U/ml hyaluronidase and observed sustained increased calcium flux of Jurkat CAR T-cells (Fig. 4B). Fig. 4C shows significantly decreased calcium flux fluctuations of CAR T-cells interacting with hyaluronidase-treated LS-174T monolayer comparable to the trypsin treated condition.

### Hyaluronidase-mediated glycocalyx degradation unmasks CEACAM5 in human colorectal cancer tissue

To determine whether the limited efficacy of anti-CEACAM5 CAR T-cell therapy in CRC patients is due to restricted antigen accessibility, we assessed CEACAM5 availability in human CRC tissue sections before and after enzymatic treatment. Immunofluorescence staining of untreated CRC tissue from multiple patients (n=13) revealed that CEACAM5 was largely inaccessible, with only sparse and heterogeneous surface expression observed across tumour regions (Fig. 5A, untreated). Control experiments using secondary antibody alone confirmed the specificity of the CEACAM5 signal and established the baseline for detection (Fig. S5A).

**Fig. 5:**
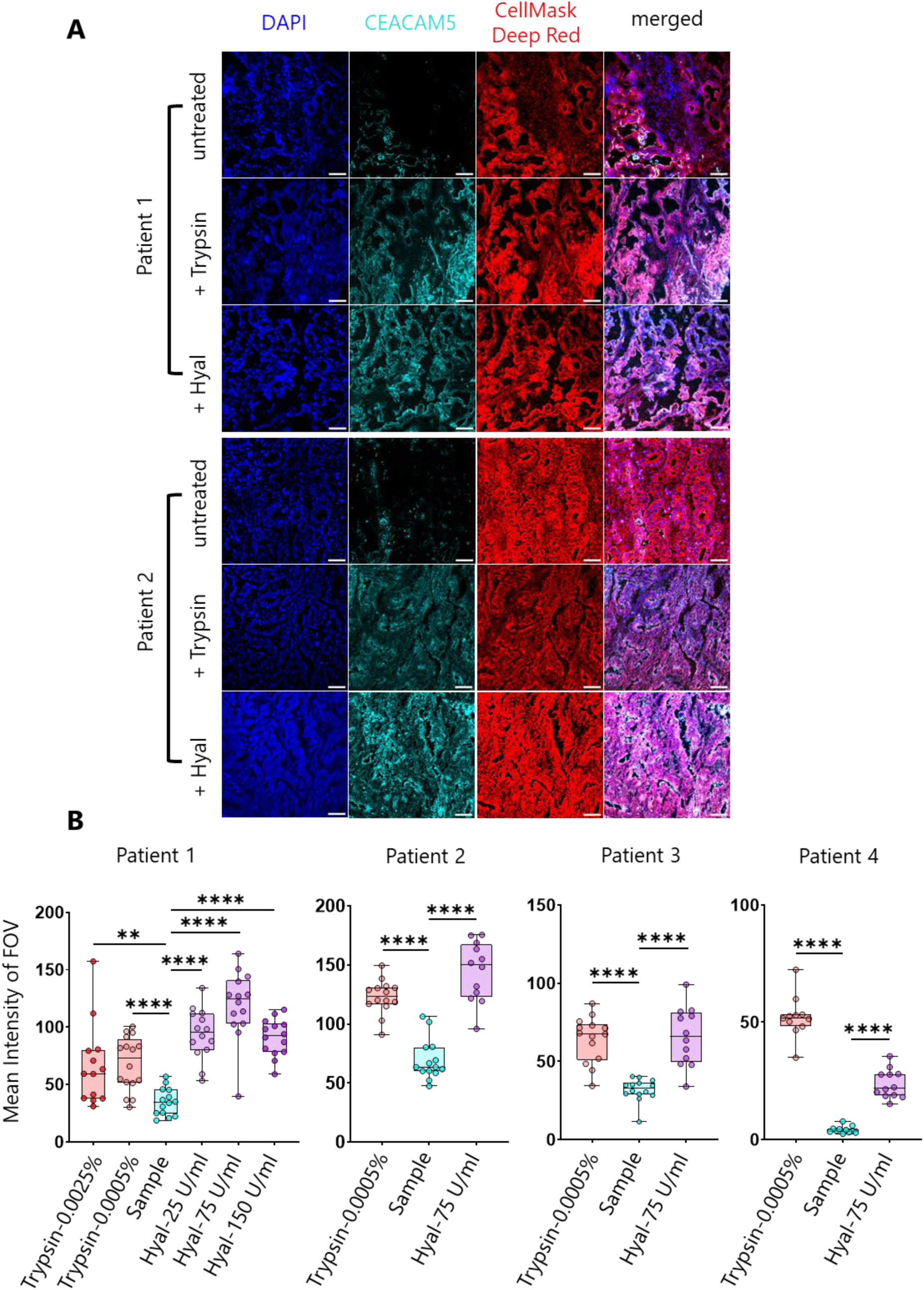
Trypsin and hyaluronidase treatment on human colorectal cancer tissue samples. (A) Representative DAPI (blue), CEACAM5 (cyan), CellMask Deep Red (red) and merged images of patient 1 and 2 tissues, and after treatment of trypsin (0.0005%) and hyaluronidase (75 U/ml). Scale bars are 50 μm. (B) Boxplots show variation of mean CEACAM5 intensity in patient 1-4 tissues, and different trypsin and hyaluronidase treatment conditions. n = 11-14 FOVs were collected in each condition and each point represents a single FOV. The medians were compared using two-sided Student’s t-test. ****P < 0.0001, **P = 0.0041.

Upon treatment with hyaluronidase (75 U/mL), a marked increase in CEACAM5 accessibility was observed. Quantitative analysis across multiple fields of view demonstrated a significant 2- to 5-fold elevation in mean CEACAM5 fluorescence intensity following hyaluronidase treatment compared to untreated controls (Fig. 5B). This effect was consistent in 4 patient samples, indicating that hyaluronidase-mediated degradation of the tumour glycocalyx efficiently unmasks sequestered CEACAM5 epitopes. Dose titration experiments established 75 U/mL as the optimal concentration for maximal antigen exposure without overt tissue disruption (Fig. 5B).

For comparison, trypsin treatment (0.0005%) also substantially increased CEACAM5 accessibility, with some tissues exhibiting up to a 12-fold enhancement (Fig. 5B). However, a subset of patient samples did not respond to trypsin enzymatic treatment, which was attributed to either intrinsic CEACAM5-negativity or preserved epithelial architecture indicative of early-stage disease (Fig. S5B-C).

Collectively, these results demonstrate that the dense glycocalyx in human CRC tissues acts as a physical barrier, concealing CEACAM5 from CAR T-cell therapeutic recognition. Hyaluronidase treatment effectively degrades this barrier, revealing previously hidden CEACAM5 and supporting the rationale for combining glycocalyx-targeting strategies with CAR T-cell therapy to improve immunotherapeutic targeting of solid tumours.

## DISCUSSION

CAR T-cell therapy has revolutionised treatment for haematological malignancies but remains largely ineffective against solid tumours like CRC. Our study identifies two critical barriers to anti-CEACAM5 CAR T-cell therapy in CRC: antigen sequestration at intercellular junctions and a substantially deepened tumour cell glycocalyx. Notably, they both contribute and we cannot easily change one without changing the other in our system. By employing cutting-edge microscopy and functional analyses, we provide mechanistic insights into CAR T-cell failure against CRC and demonstrate promising enzymatic strategies to restore therapeutic efficacy.

### Antigen sequestration and glycocalyx barriers limit CAR T-cell functionality

We show for the first time that CEACAM5 becomes progressively sequestered at intercellular junctions as CRC monolayers mature, rendering this target antigen increasingly inaccessible to CAR T-cells. This sequestration directly correlates with the transition from sustained to oscillatory calcium signalling in CAR T-cells and abolished cytotoxicity against confluent tumour cell monolayers. The antigen-specific nature of this phenomenon is evidenced by our observation that other membrane proteins (CD19, HER2), which are successfully targeted by CAR T-cell therapy, remain homogeneously distributed and accessible regardless of monolayer maturity.

Super-resolution imaging revealed that CRC cells possess a remarkably deep glycocalyx, approximately four times deeper than that of CAR T-cells. This substantial physical barrier likely explains the delayed CAR T-cell activation kinetics observed in our experiments, as the receptor must overcome this steric hindrance to engage target antigens. The combination of antigen sequestration and glycocalyx barrier presents a formidable challenge for CAR T-cell efficacy against solid tumours.

### CD45 exclusion supports the kinetic-segregation model in CAR T-cell activation

We found that CD45 exclusion is a potential mechanism of CAR T-cell activation at the anti-CEACAM5 CAR T-cell/CRC cell interface. Along with other studies, our work confirms and extends the reach of the kinetic segregation (KS) model, wherein CD45 exclusion initiates receptor triggering, to CAR T-cells at T-cell/tumour interfaces irrespective of the tumour types (liquid and solid cancer) (*14, 15*).

### Local and bulk treatments for enzymatic remodelling

To achieve precise intercellular dissociation without compromising the monolayer of cells to avoid glass/T-cell interaction, we employed local trypsin treatment using a micropipette (Fig. 4B, 4C). This treatment was also used to investigate the CEACAM5 re-availability kinetics (Fig. 4D, S4C), ensuring same FOV before and after trypsin treatment. However, to simplify the experimental procedure, bulk treatment was employed during resPAINT measurement (Fig. 4E) and experimenting with hyaluronidase (Fig. 4B, 4C). Our results demonstrated both local and bulk treatments are working.

### Enzymatic remodelling restores CEACAM5 accessibility and CAR T-cell function

Our most significant finding is that targeted enzymatic treatments effectively overcome both identified barriers. Localised trypsin application disrupted intercellular junctions, mobilised sequestered CEACAM5, and significantly reduced glycocalyx thickness. This dual mechanism of action restored sustained calcium signalling in CAR T-cells and reinstated their cytotoxic capacity against previously resistant CRC monolayers.

Importantly, hyaluronidase treatment, which specifically targets the glycocalyx without disrupting cell-cell junctions, similarly enhanced CAR T-cell activation. This finding has immediate clinical relevance given hyaluronidase’s established safety profile and FDA approval for other indications. The ability of two mechanistically distinct enzymes to improve CAR T-cell function provides compelling evidence that physical barriers, rather than intrinsic tumour cell resistance mechanisms, are primary impediments to CAR T-cell efficacy in CRC.

### Pre-clinical validation and therapeutic implications

Extending beyond in vitro models, we demonstrated that CEACAM5 is largely inaccessible in patient-derived CRC tissues, with enzymatic treatment significantly enhancing antigen detection. The differential response rates between hyaluronidase and trypsin treatment highlight tumour heterogeneity and suggest that glycocalyx remodelling may have broader applicability than junctional disruption for clinical translation.

Our findings provide a mechanistic explanation for the paradoxical clinical observation of respiratory toxicity despite limited efficacy in early CEACAM5 CAR T-cell trials (*16*). If CEACAM5 is sequestered in colorectal tumours but remains accessible on normal lung epithelium, CAR T-cells would preferentially target non-malignant tissue, explaining the observed on-target, off-tumour effects.

### Integration with emerging approaches to enhance CAR T-cell therapy

This work complements recent advances in solid tumour immunotherapy strategies (*25–27*). While some approaches focus on enhancing CAR T-cell trafficking or persistence, our findings address the fundamental barrier of antigen accessibility at the tumour-immune interface. Our results align with studies demonstrating that disrupting tumour glycosylation enhances immunotherapy efficacy (*25*), but provide a potentially more clinically feasible approach through enzymatic preconditioning rather than genetic manipulation of glycosylation pathways. Zhao et al. (*27*) demonstrated hyaluronidase degrades hyaluronic acid enabling CAR T cells to infiltrate deeply into solid tumours. We believe hyaluronidase/trypsin degrades the glycocalyx and unmasks sequestered CEACAM5 in longer-seeded monolayer/cancer tissues.

Future enhancement with the spatial control afforded by localised enzymatic delivery presents advantages over systemic approaches, potentially minimizing off-target effects while maximising tumour impact. Furthermore, our findings suggest that combinatorial strategies, pairing CAR T-cells with tumour microenvironment-modifying agents, may be necessary to achieve optimal therapeutic outcomes in solid tumours.

### Limitations and future directions

While our study provides compelling mechanistic insights, some limitations must be addressed in future work. First, the in vitro and ex vivo nature of our experiments necessitates validation in immunocompetent in vivo models. Second, clinical translation requires development of targeted enzyme delivery strategies to avoid systemic effects. Third, the long-term impact of glycocalyx remodelling on tumour biology requires careful evaluation. Fourth, an experiment where anti-CD19 CAR T-cells interact with CD19 expressing SLB2 (40 nm glycocalyx) and CD19^+^ LS-174T (400 nm glycocalyx) monolayer will be helpful to evaluate the effect of the glycocalyx alone on CAR T-cell triggering.

Future studies should explore the following promising directions: (1) development of enzyme-armed CAR T-cells capable of autonomously modifying their local microenvironment; (2) optimisation of hyaluronidase dosing and scheduling to maximise the therapeutic window; (3) exploration of combinatorial approaches with checkpoint inhibitors or stroma-targeting agents; and (4) identification of biomarkers to stratify patients likely to benefit from this approach.

### Conclusion

Our study reveals antigen sequestration and glycocalyx barriers as key mechanisms limiting CAR T-cell efficacy in CRC and demonstrates that enzymatic remodelling of the tumour microenvironment can effectively overcome these barriers. By elucidating these fundamental limitations and providing a rationalized approach to address them, this work establishes a framework for improving immunotherapy outcomes in colorectal cancer and potentially other solid malignancies. The use of clinically approved agents like hyaluronidase offers a pathway for rapid translation of these findings to benefit patients with currently incurable solid tumours.

## METHODS

### Isolation and culture of human primary CD8^+^ T-cells

Fresh peripheral blood was obtained from three healthy donors after informed consent. Peripheral blood mononuclear cells (PBMCs) were enriched by density gradient centrifugation using Ficoll-Paque Plus (GE Healthcare). Naïve CD8 T-cells were isolated from PBMCs using the MagniSort Human CD8 Naïve T-cell enrichment kit according to the manufacturer’s instructions (Invitrogen). 1 million naïve CD8^+^ T-cells were activated with plate-bound anti-CD3 (Biolegend Ultra-LEAF purified, Clone-OKT3, at 1 μg/mL) and anti-CD28 (Biolegend Ultra-LEAF purified, Clone-CD28.2, at 1 μg/mL) for 3 days. Primary CD8^+^ T-cells were cultured in Immunocult (Stem Cell Technologies) medium supplemented with 50 μg/mL of Gentamicin (Gibco) and 50 U/mL of human recombinant IL-2 (Peprotech). IL-2 was replenished after every 3 days.

### Generation of CAR T-cells and overexpression of human CD19 by Sleeping Beauty (SB)

Sleeping beauty (SB) transgene cassettes were used to facilitate overexpression of human CD19 and expression of second-generation CARs targeting human CEACAM5 and CD19. All parental plasmids (pp) were designed and purchased from VectorBuilder. The CD19 SB-CAR was constructed using NEBuilder® HiFi DNA Assembly Master Mix (New England Biolabs, #E2621S) according to the manufacturer’s instructions. The two assembled fragments were PCR-linearized CEACAM-hMN-14 SB-CAR (with the scFv removed) and PCR-amplified anti-CD19 FMC63 single chain variable fragment (scFv) in the VL-Whitlow/218 linker-VH orientation. For CD19 overexpression in LS-174T, a puromycin resistance gene was incorporated into the transposon cassette. For anti-CEACAM5 CAR expression in primary CD8 T-cells, the hMN-14 SB vector was synthesized into a pre-clinical grade (pGC) Gencircle (Genscript Biotech). Stable expression in cell lines was facilitated by SB100X (VectorBuilder), while SB100X mRNA (N1-Methylpseudouridine/m1Ψ) (Genscript Biotech) was employed for primary CD8^+^ T-cells. All plasmid transgene cassettes except for SB100X, which uses the CAG promoter, were under control of EF1a.

### CAR T Structure

The anti-CEACAM5 scFv comprised variable domains (VH and VL) derived from the clone hMN-14 with FMC63 being used for CD19. In contrast, to anti-CD19 the CEACAM5 variable domains were linked by a poly-Glycine-Serine (G4S)3 linker. Both CAR constructs incorporated a Strep-tag II (WSHPQFEK) sequence and a G4S2 linker, fused between the CD8α hinge and scFv domains to facilitate identification of CAR^+^ cells. The transmembrane domain from CD28 was used in conjunction with the intracellular signaling domains of CD28 and CD3ζ.

### Nucleofection and selection of SB delivered cassettes

All electroporations were carried out using a Lonza 4D Nucleofector. Briefly, 1-2 million cells were resuspended in their corresponding nucleofection buffer and pulsed according to their respective code (see the following table). Following nucleofection, cells were immediately rescued with 80 μL of pre-warmed complete media (antibiotic-free) and incubated at 37°C for 20 minutes. Subsequently, cells were further diluted and cultured in complete media. LS-174T cells were selected for CD19 expression by the addition of puromycin (1 μg/mL) (Santa-Cruz) for one week. Jurkat CAR^+^ cells (anti-CD19 & CEACAM5) were stained with anti-strep-tag II antibody (FITC) and enriched using the EasySep FITC Positive Selection Kit II according to the manufacturer’s instructions (StemCell Technologies). Conversely, 72-hours post-nucleofection CD8^+^ CAR T-cells were expanded with plate-bound CEACAM5 protein with FC tag (ACROBiosystems) at 0.5 μg/mL for 4 days in complete Immunocult media with 50 U/mL of human IL-2. The percentage of CD8^+^ CAR^+^ cells was subsequently determined by flow cytometry.

| Cell Type/Line | No. of Cells | Buffer Solution | Pulse Code |
| --- | --- | --- | --- |
| Jurkat | 1 million | SE | CL-120 |
| LS-174T | 1 million | SE | EN-113 |
| Human CD8<sup>+</sup> T-cells | 2 million | P3 | EO-115 |

### Flow Cytometry

Flow cytometry was performed on a CytoFLEX LX Flow Cytometer, with all analysis done on FlowJo software (v10.10). Primary and CRC cell lines were stained for viability with LIVE/DEAD Fixable Near-IR and blocked with Human TruStain FcX (Biolegend) prior to antibody staining. For antibody staining, cells were resuspended in FACS buffer (PBS with 3% FBS). The following antibodies were used: anti-human CD8 Pacific Blue, (Biolegend, RPA-T4), The NWSHPQFEK Tag Antibody FITC, (Genscript, 5A9F9), anti-human CD19 AF594 (Biolegend, HIB19) anti-human CD340/HER-2 AF647 (Biolegend, 24D2) and anti-human CEACAM5/CD66e PE (R&D Systems, 487609). Cells were fixed in IC Fixation Buffer (eBioscience) and washed in FACS buffer twice prior to acquisition.

### Jurkat CAR T-cell and CRC cell culture

Jurkat CAR T-cells (CEACAM5^+^, CD19^+^) and CRC cells (CEACAM5^+^ & HER2^+^ WT LS-174T, CEACAM5^-^ RKO, CD19^+^ LS-174T, CEACAM5^+^ LS-1034) were cultured in StableCell^TM^ RPMI-1640 and DMEM/F-12 (1:1) (1X) + GlutaMAX^TM^-I (Dulbecco’s Modifies Eagle Medium F-12 Nutrient Mixture (Ham)) medium, respectively. Both media was supplemented with 10 % (v/v) fetal calf serum (FCS), 1 % (v/v) HEPES buffer solution (1M), 1 % (v/v) sodium pyruvate 100 mM (100X) and 1 % (v/v) pen strep antibiotics. Hereafter, we will call the supplemented media as ‘complete RPMI’ and ‘complete DMEM’. TrypLE^TM^ express enzyme was used to detach the CRC cells. Cells were then diluted and re-suspended in complete DMEM for further culture. CRC cells were passaged before they could grow confluent. All cell culture was performed in HEPA-filtered cell culture cabinets and cells were grown within the incubator with 37^0^C and 5 % CO_2_ maintaining density 0.2-0.5 million/ml.

### SLB2 preparation and monitoring CAR T-cell/SLB2 interaction using imaging

A mixture of 98:2 mol% POPC (Avanti Polar Lipids) and DGS-NTA-Ni^2+^ (NiNTA; Avanti Polar Lipids) was prepared as previously described for making SLB2 using the vesicle fusion technique (*17*). SLB2 protein mixture presents CD58 and ICAM-1, CD45 and CD43, human CEACAM5/CD66e protein (from ACROBiosystems), and gp100-MHC. CD45 and CD43 proteins have been conjugated covalently with Alexa-555 using random lysine before use. CEACAM5 targeting Jurkat CAR T-cells were labelled with a mixture of Fluo-4-AM (calcium indicator) and Cell-Mask Deep Red Plasma Membrane Stain (ThermoFisher) before being allowed to interact with SLB2. As experimental readouts, calcium signaling, cell footprint, and close-contact (CD45/CD43 exclusion) formation were measured using a home-built microscope. Individual protein concentrations of CEACAM5^+^ and CEACAM5^-^ SLB2s are given below:

|  | CEACAM5<sup>+</sup>SLB2 | CEACAM5<sup>-</sup>SLB2 |
| --- | --- | --- |
| Protein | Conc. (ng/μL) |  |
| CEACAM5 | 0.5 | 0 |
| gp100-MHC | 9.5 | 10 |
| CD58 | 0.5 | 0.5 |
| ICAM-1 | 1.2 | 1.2 |
| CD45 | 2.5 | 2.5 |
| CD43 | 1.5 | 1.5 |

Details of protein production, labelling, SLB2 preparation and experiments have been described in our previous publication (*17*).

### TIRF Microscopy

The interaction of CAR T-cell/SLB2 was imaged using a home-built microscope in total internal reflection fluorescence (TIRF) mode with three laser lines: 488 nm (200 mW, iBeam-SMART, Toptica), 561 nm (LPX-561L, LaserBoxx, Oxxius), and 641 nm (Obis, Coherent). Each laser line was attenuated by neutral density filters, expanded by a pair of plano-convex lenses, circularly polarised by a quarter-wave plate, and combined in the downstream path by appropriate dichroic mirrors. The lasers were then aligned and directed to the edge of a 100X oil-immersion objective lens (CFI Apo TIRF 100XC Oil, NA 1.49, MRD01991, Nikon), which was mounted on an inverted optical microscope (Eclipse Ti2, Nikon). Fluorescence emission was collected through the same objective lens, and a quad-band dichroic mirror (Di01-R405/488/561/635, Semrock) was used to separate the emission from the excitation light. The built-in 1.5X magnifying lens in the Ti2 microscope body was used to further improve the image resolution. Emission was then filtered through appropriate long-pass and single-band filters (BLP01-488R + FF01-520/44-25, LP02-568RS-25 + FF01-587/35-25, FF01-692/40-25; Semrock) before being collected by an EMCCD camera (Evolve 512 Delta, Photometrics). The microscope was equipped within an incubator (DigitalPixel) to maintain 37 °C during live T-cell imaging. The built-in Perfect Focus System (PFS) of the TI2 microscope body was used to maintain focus during imaging, and three-colour images were acquired sequentially using Micro-Manager 2.0.0 software with a 100 ms exposure time and a 2-second interval. The pixel size of the collected images was 107 nm, measured using a line grating target (R1L3S6P, Thorlabs).

### CAR T-cell/SLB2 imaging data analysis

Data was analysed using a custom-written code described in our previous publication (*17*).

### CRC cells’ (WT LS-174T, LS-1034, RKO, CD19^+^ LS-174T) monolayer preparation

The viability of the adherent CRC cells was checked and counted using Countess II automated cell counter (ThermoFisher) by mixing with 1:1 trypan blue (0.4%). The cells were plated on ibidi glass bottom (35 mm, high) dish / 8 well plate maintaining 0.3-0.6 million/ml and incubated for seeding for 24 and 48 hrs (early and longer, respectively).

### Calcium imaging and analysis

On the day of the experiment, CAR T-cells were labelled with Fluo-4-AM and allowed to interact with CRC cells’ monolayer. As previously mentioned, a 488 nm laser was used for excitation, and epifluorescence calcium imaging of CAR T-cells was monitored at the T-cell/tumour interface. Imaging was performed with an exposure time of 100 ms with a 30-second interval. Initially, a 20X objective lens (Plan Fluor, NA 0.50, Nikon) was used to capture a large field of view (FOV) of T-cell/tumour interactions. Subsequently, a 60X oil-immersion objective lens (CFI Apo TIRF 60XC Oil, NA 1.49 oil-immersion, MRD01691, Nikon) was used for close inspection. The pixel size of the collected images using 20X and 60X objective lens were 800 nm and 178 nm, respectively.

Calcium imaging time-stacks were analysed using the TrackMate plugin in ImageJ/Fiji (*28*). Cells within the imaging FOVs were detected and segmented using the StarDist (*29*) detector in TrackMate with default parameters. These segmented cells were linked into trajectories using the simple LAP tracker to capture the dynamics of a T-cell in all frames throughout the movie. Parameters for linking max distance, gap-close max distance and gap-close max frame gap were set to ensure that segmented cells from the same cell across different frames were accurately joined. Typical parameters were 15 μm for max distances and 10 frames for max frame gap. TrackMate outputted a list of polygon regions of interest (ROI) for the segmented cells, with each ROI labelled according to the track it belonged to. The mean fluorescence intensity of each cell across all frames was measured using the Measure function in ImageJ/Fiji (Fig. S5D). The results and the ROI information were saved to local files and organized using a home-written Python script, which processed the data to output intensity information as a function of time for each individual cell. Mean fluorescence intensity (I_0_ indicates the first frame where T cell activity is detected; I_t_ denotes the subsequent frames over time) was extracted and the ratio (I_t_/I_0_) was plotted as a function of time for each cell using OriginPro 2023b.

For comprehensive comparison of intensity fluctuations of Jurkat and primary CAR T-cells in triggering/non-triggering conditions, the intensity profile of each tracked cell was normalised to its maximum intensity. The variance of each profile was then computed and recorded. Violin plots were generated using GraphPad Prism 10.3.0 to visualise the distribution of intensity variations.

### CAR accumulation experiment

FITC conjugated anti-strep-tag II antibody was used to label the strep-tag II of the CEACAM5^+^ Jurkat CAR T-cells and they were allowed to interact with early-seeded LS-174T monolayer. A 488 nm laser with 100 ms exposure time was used as excitation source and CAR accumulation was monitored by epifluorescence imaging with time.

### Glycocalyx measurement of T-cell and LS-174T using super-resolution imaging

CAR T-cells and early-, longer-seeded LS-174T cells were fixed using a mixture of 0.8 % paraformaldehyde (28906, Thermo Scientific, MA) and 0.2 % glutaraldehyde (G5882, Sigma-Aldrich) at room temperature. Fixed CAR T-cells were allowed to settle on the poly-L-lysine (PLL, 150-300 kDa; P4832; Sigma-Aldrich) coated coverslip for 45 min as described in our earlier publication (*24*). Sodium carbonate-bicarbonate buffer with pH 9.6 was used to dilute HMSiR conjugated wheat germ agglutin (WGA) and used to label the glycocalyx of the cells. A 641 nm laser in highly inclined and laminated optical sheet (HILO) mode was used to excite the sample and 20000 frames were collected with 30 ms exposure time. 2D resPAINT super-resolution imaging data was analysed in Fiji using PeakFit (GDSC SMLM 2.0) and shown in Fig. 2 (*24*).

Diluted 0.05% Trypsin-EDTA (1X, gibco) was used for bulk treatment of early- and longer-seeded LS-174T monolayers. Trypsin treated monolayers were washed and fixed as described before. resPAINT imaging was carried out and the results are shown in Fig. 4E.

### Antigen mapping on LS-174T monolayer using antibody

PE conjugated anti-hCEACAM5 (mouse IgG_2A_, R & D Systems), Alexa Fluor 594 conjugated anti-hCD19 (mouse IgG_1_, R & D Systems) and Alexa Fluor 647 conjugated anti-human CD340 (erbB2/HER-2, Biolegend) antibodies were used for CEACAM5, CD19 and HER2 mapping, respectively. Respective lasers were used for excitation and epifluorescence images were collected.

### Granules release experiment and analysis using Imaris

Primary CAR T-cells were labelled with a mixture of Fluo-4-AM and lysotracker red (50 nM) and allowed to interact with early-seeded LS-174T monolayer. 488 nm and 561 nm lasers were used for excitation and epifluorescence images were collected to monitor calcium flux and granule release by CAR T-cells simultaneously (Fig. 3C, 3D).

### Granules release analysis using Imaris

Released granules from individual T-cells were tracked using Imaris 10.0.1 (Oxford Instruments). The surface function with tracking was utilized for detecting the granules. This tool allows us to create an artificial solid object (surfaces) to represent the range of interest of a specific object, like granules here. The automatic surface creation wizard in Imaris was used for segmenting granules, followed by fine-tuning the segmentation for accuracy. We used in-built tools in Imaris for this purpose, which involves adjusting the contour of the surface or correcting any segmentation errors. To make the surfaces more visible, we assigned different colours and opacity to the created surfaces. We then tracked the resulting surfaces over time using an autoregressive motion tracking algorithm. Several parameters, such as track length, track speed mean, and track straightness, were computed.

#### Track length

The Track Length (*tl*) is the total length of displacements within the Track.

#### Track speed mean

The mean value of the cell’s speed on the track. Mean speed is determined by dividing track length by the time between first and last object in the track.

#### Track straightness

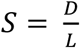

S = Track Straightness

D = Track Displacement

L = Track Length

### Segmenting and Tracking CAR T-cells using Imaris

Calcium imaging movies of Jurkat CAR T-cell/LS-174T interaction were monitored using 60X objective lens and used for analyzing track speed mean and track straightness of CAR T-cells using Imaris as described in previous section.

### Imaging CD45 exclusion at T-cell/tumour cell interface using eSPIM

#### Optical Setup

The experiment employed an open-top single-objective light sheet microscope based on the Oblique Plane Microscopy (OPM) configuration (*30*). The instrument adopted a similar design as reported by Yang et al (*31*). A primary 60X NA 1.27 water-immersion objective (CFI Plan Apo IR 60XC WI, Nikon) mounted on an inverted microscope body (Eclipse Ti-U, Nikon) was used for both light sheet launching and emission collection. Two continuous diode lasers, 488 nm (200 mW, LBX-488-200, Oxxius) and 638 nm (180 mW, 06-MLD-638, Cobolt), were expanded, collimated and combined into a single excitation path. The combined beam then passed through a cylindrical lens to generate a 1D Gaussian beam. A quad-band dichroic mirror (Di01-R406/488/561/635, Semrock) was used to separate the excitation and emission wavelengths. The beam propagated through two 4f systems before entering the primary objective at one edge of the back focal plane (BFP) to produce a 30-degree-tilted light sheet. Rapid volumetric illumination was achieved by scanning the light sheet using a 1-D Galvo mirror (GVS201, Thorlabs). The operation of the galvo mirror was controlled by a home-written LabView script.

Fluorescence collected by the primary objective was transmitted through the quad-band dichroic mirror, relayed onto a 40X secondary air objective (CFI Plan Apo Lambda D Air, NA 0.95, Nikon), with the pupil plane conjugated to that of the primary objective. Subsequently, the emission was collected by a bespoke glass-tipped tertiary objective (AMS-AGY v1.0, NA 1, Calico lab) oriented at a 30-degree angle to the optical axis. A tube lens (F = 321 mm) was used to direct the emission onto a sCMOS camera (Photometrics Prime 95B). Additionally, the setup featured a three-colour optical splitter (OptoSplit III, Prior) for simultaneous multi-colour imaging.

### Experimental Details

LS-174T and Jurkat CAR T-cells were labelled with Alexa Fluor 647 anti-human CD326 (Ep-CAM, clone 9C4, BioLegend) antibody and Alexa Fluor 488 anti-CD45 monoclonal antibody (Gap8.3), respectively and CAR T-cells were allowed for interaction with early seeded LS-174T monolayer and incubated for 30 mins to image late-stage contact formation. Volumetric scans were performed using a home-written automation script in μManager 2.0. During each volumetric scan, the light sheets from the 488 nm (44.4 W/cm^2^) and 638 nm (55.5 W/cm^2^) lasers scanned through the sample. The detection camera was synchronised with the galvo scanning, enabling simultaneous three-colour imaging with an exposure time of 30 ms. Each volumetric scan consisted of 100 frames (full 1200 x 1200-pixel FOV) acquired within approximately 4 seconds.

To monitor the late-stage dynamics at CAR T-cell/LS-174T cell interface, volumetric scans were repeated every 30 seconds for approximately 40-45 minutes. Throughout the acquisition process, the sample stage and the primary objective were enclosed within a sealed plastic chamber. The temperature around the sample was maintained at 37°C. The acquired images from each volumetric scan were deskewed and reconstructed into 3D projections using a custom Python script.

### Micropipette preparation and trypsin delivery

#### Micropipette fabrication

Micropipettes were fabricated from borosilicate glass capillaries (World Precision Instruments, 1B150F-4), with an outer and inner diameter of 1.5 and 0.84 mm, respectively. A laser pipette puller was used for fabrication (Model P-2000, Sutter Instrument, CA) with the following parameters: (Line 1) Heat = 415, FIL = 4, VEL = 30, DEL = 200. This pulled a micropipette with a pore diameter of approximately 4-5 µm (Fig. S5E).

#### Micropipette delivery

The micropipette was controlled using a 3D manipulator (Scientifica PatchStar Micromanipulator, Scientifica, Uckfield, UK) to provide precise positioning of the tip above the cell. A continuous pressure pulse (0.7 psi or 5 kPa) was used to deliver trypsin, until clear disruption of the cell contacts in the 2^nd^ layer was observed. Careful control of the pressure pulse ensured that the cell monolayer was not disturbed in this process. A similar strategy was employed for local delivery of the fluorescent antibody.

#### Collection of human colorectal cancer tissue

Human samples utilized in this research project were collected from the Imperial College Healthcare Tissue Bank (ICHTB), which is supported by the National Institute for Health Research (NIHR) Biomedical Research Centre at Imperial College Healthcare NHS Trust and Imperial College London. The ICHTB, approved by Wales REC3 (22/WA/0214) for releasing human material for research, issued the samples under sub-collection reference numbers associated with project R24029.

#### Immunofluorescence staining of frozen colorectal cancer tissue sections

Optimal cutting temperature (OCT)-embedded frozen human colorectal cancer samples were sectioned and mounted on glass slides. Tissue sections were treated with phosphate-buffered saline (PBS) (for untreated condition) or a specific concentration of trypsin or hyaluronidase for 15 minutes at room temperature. Following enzymatic digestion, slides were rinsed with PBS and fixed with 1% formalin solution (Sigma-Aldrich) in PBS for 10 minutes at room temperature. After fixation, slides were washed three times with wash buffer (PBS supplemented with 0.5% Tween-20) and permeabilized with 0.2% Triton X-100 in PBS for 10 minutes. This was followed by three additional washes with wash buffer. To block non-specific binding, sections were incubated with 10% goat serum (Invitrogen 31873) in wash buffer for 1 hour at room temperature. Slides were then incubated overnight at 4 °C in a humidified chamber with a mouse monoclonal anti-human CEACAM5 primary antibody (CB30, S-52390; 1:500 dilution) prepared in 5% goat serum in wash buffer. The following day, slides were washed three times with wash buffer (10 minutes each wash at room temperature), followed by a 1-hour incubation at room temperature in the dark with Alexa Fluor 488-conjugated goat anti-mouse IgG secondary antibody (Invitrogen; 1:2000 dilution in 5% goat serum in wash buffer). After secondary antibody incubation, sections were washed three times with wash buffer and counterstained with DAPI (1:2000 dilution) for 10 minutes. Membranes were then stained with CellMask Deep Red according to the manufacturer’s protocol. Final washes were performed three times in wash buffer before slides were mounted using CitiFluor AF1 mounting medium.

### Tissue imaging

Imaging was performed using a confocal microscope (STELLARIS8, Leica, Wetzlar, Germany). Images were acquired using a frame size of 512 x 512 pixels in the violet (405 nm excitation, 420-470 nm emission), green (488 nm excitation, 495-545 nm emission), and the far-red (637 nm excitation, 645-700 nm emission) channel. In a FOV, upper and lower Z-positions were selected based on the emission signal from the green channel (CEACAM5) and images were acquired as Z-stake with 0.5 µm inter-stake separation.

### Tissue imaging analysis

The tissue image stack consists of images captured at different axial depths. To quantify the level of exposed CEACAM5 before and after trypsin treatment, we compared the mean intensity of the FOV before and after treatment. Image analysis was performed in Fiji/ImageJ. Each Z-stack was first processed using a maximum intensity Z-projection to generate a single representative image of the entire FOV. The resulting projection was then converted to 8-bit, and an intensity threshold of 20 to 255 was applied. The mean intensity of the FOV was measured using the ’Measure’ function in ImageJ and recorded. The same analysis and intensity threshold were consistently applied to images from all patient samples. Multiple FOVs across 13 patient samples were analysed. The mean FOV intensity before and after trypsin treatment was visualised using box plots for each patient sample.

### Statistical analysis

Statistical analysis was performed using Graphpad Prism 10.3.0. Two-sided paired and unpaired Student’s t-test were performed as required for statistical significance in Fig. 1-5 and Fig. S3, S5.

## Data and code availability

All imaging data needed to evaluate the conclusion are present in the paper and the supplementary materials and movies. Codes are available in Github (https://github.com/mkoerbel/contactanalysis_2D, https://github.com/Zui409 & https://github.com/Zui409/Intensity-tracking/tree/main). All additional datasets of the current study are available from the corresponding authors on request.

## Acknowledgments

We gratefully acknowledge the flow cytometry facility from the School of the Biological Sciences for their support and assistance, and Cambridge Advanced Imaging Centre (CAIC) for providing access to their data analysis workstation and softwares. Additionally, we would also like to give thanks to Genscript Biotech for kindly providing pre- clinical grade Gencircles and SB100X mRNA (N1-Methylpseudouridine/m1Ψ).

## Author contributions

Conceptualization: D.B. and D.K. Methodology: D.B., P.S., B.L., D.H., S.K., Z.Z., S.J.D., and D.K. Cells and reagents provided: C.W., J.P.R., M.A.C., S.J.D., and B.M. Investigation: D.B., C.W., S.C., P.S., B.L., and D.H. Visualization: D.B., C.W., S.K., and Z.Z. Funding acquisition: D.B., B.M., S.J.D., and D.K. Project administration: D.B. and D.K. Supervision: D.K. Writing-original draft: D.B. and D.K. Writing-review & editing: D.B., S.J.D., B.M., D.K.

## Additional information

Supplementary material contains Fig. S1-S5, figure legends and description of supplemental movies S1 to S15.

**Correspondence** and requests for materials should be addressed to Debasis Banik, Bidesh Mahata or David Klenerman.

