## Supplementary materials for "Hyaluronidase unlocks sequestered CEACAM5 and improves CAR T-cell therapy in colorectal cancer"

**Supplemental Figures**

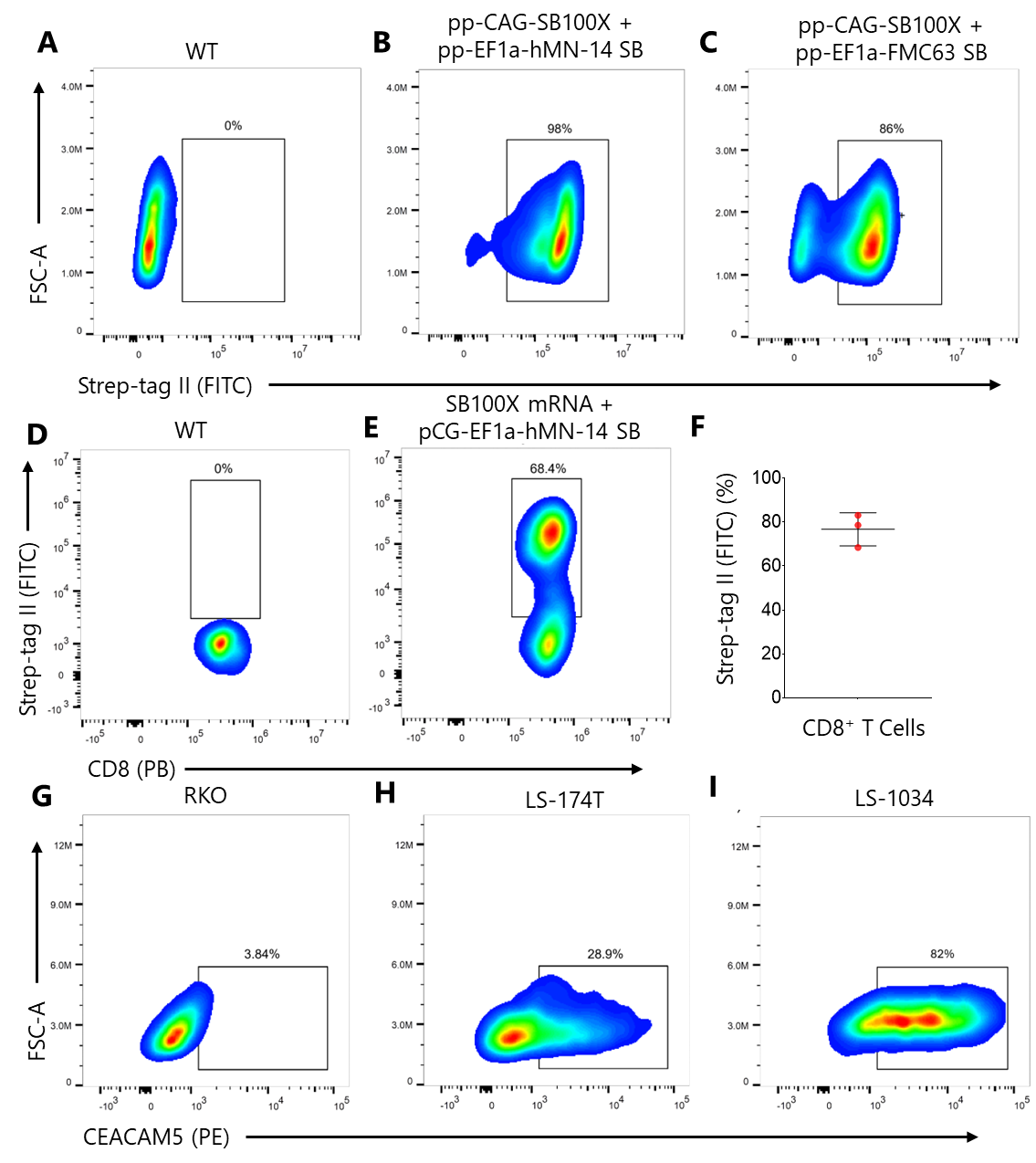

Fig. S1: (A-C) Representative flow cytometry data of CAR expression (Strep-tag II) in Jurkat. (A) CAR expression in WT Jurkat cells (control). 1 million Jurkat cells were electroporated with 250 ng of pp-CAG-SB100X and (B) 1 µg of pp-EF1a-hMN-14 SB or (C) pp-EF1a-FMC63 SB. 1 week after nucleofection, cells were stained with anti-strep-tag II antibody (FITC) and enriched using a FITC positive selection kit (Easy Sep^TM^). CAR expression was determined 72 hours later by flow cytometry.

(D-F) Representative flow cytometry data of CAR expression (Strep-tag II) in human CD8 T-cells. (D) WT CD8^+^ T-cells without CAR expression. (E) 2 million CD8^+^ T-cells were electroporated with 4.8 µg of SB100X mRNA and 1.2 µg of pGC-EF1a-hMN-14 SB. 72-hours post nucleofection, 1 million CD8^+^ T-cells were seeded onto 24-well plates containing 0.5 µg/mL of CEACAM5 (Fc tag) protein. After 4 days of stimulation, CAR expression was determined by flow cytometry. (F) Variation in CAR expression in CD8^+^ T-cells. Anti-Strep II tag antibody was used to evaluate hMN-14 CAR expression in human CD8^+^ T-cells after 4 days of enrichment with CEACAM5 (Fc tag) protein (n = 3). Error bar depicts Mean ± SD.

(G-I) Representative flow cytometry data of CEACAM5 expression in colorectal cell lines. (G) RKO, (H) LS-174T and (I) LS-1034.

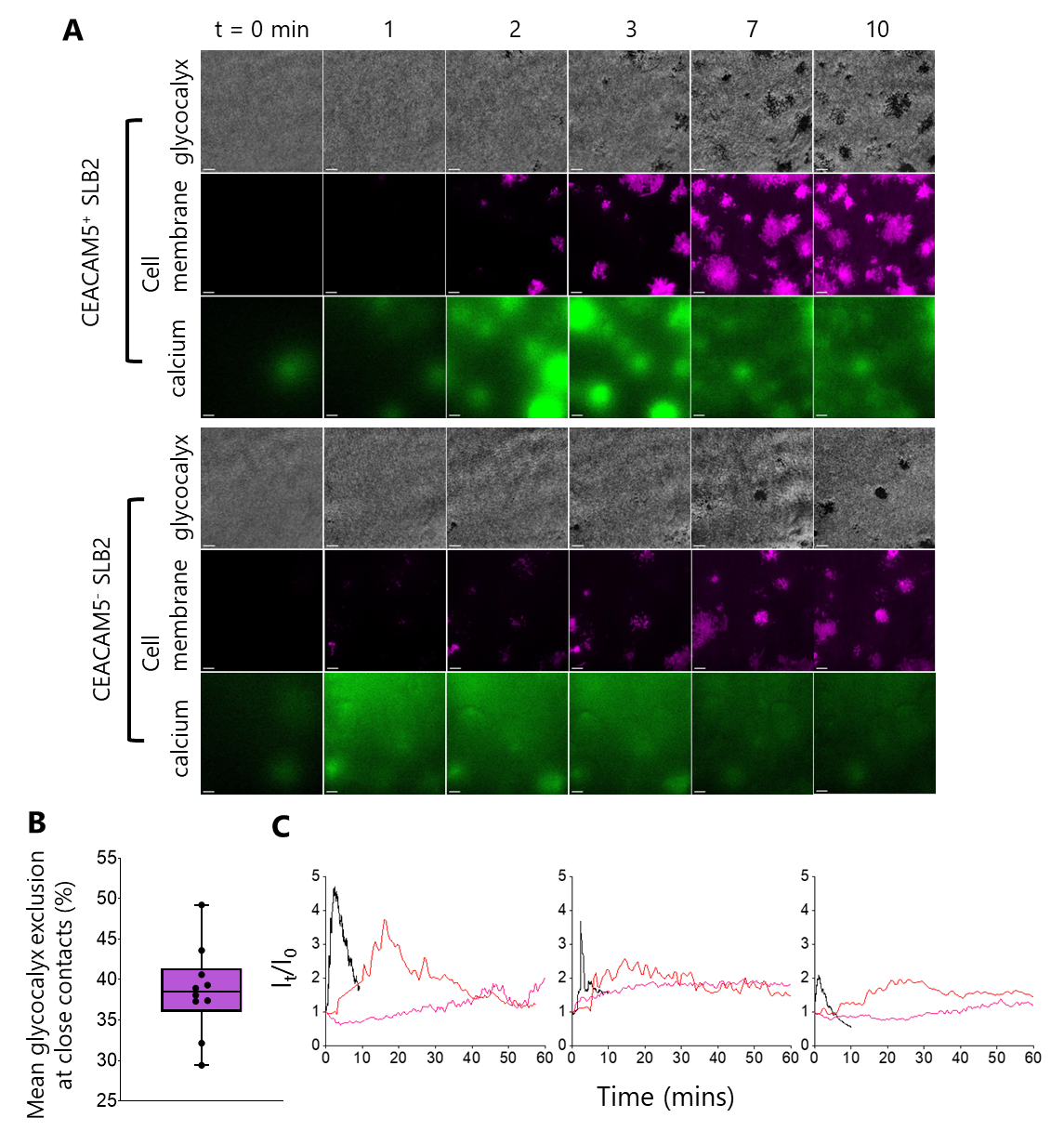

Fig. S2: (A) Time-series showing the interaction between anti-CEACAM5 Jurkat CAR T-cells and CEACAM5^+^ and CEACAM5^-^ SLB2s.

(B) Box plot shows mean glycocalyx (CD45/CD43) exclusion at close contacts formed prior to calcium signaling while CAR Jurkat T-cells interacting with CEACAM5+ SLB2. Data points were extracted by analyzing ten (n = 10) T-cells from three independent experiments. Each data point represents the average glycocalyx exclusion across all close contacts formed by one cell.

(C) Comparison of calcium flux profile of CAR Jurkat T-cells on CEACAM5+ SLB2 (black), early seeded LS-174T monolayer (pink), trypsin treated early seeded LS-174T monolayer (red). While instant triggering was observed on SLB2 (peak maximum appeared between 1-3 mins), delayed triggering was found on LS-174T monolayer (in most cases, we didn't see peak maximum within one hour of monitoring). In presence of trypsin, triggering was faster and peak maximum appeared within 30 mins.

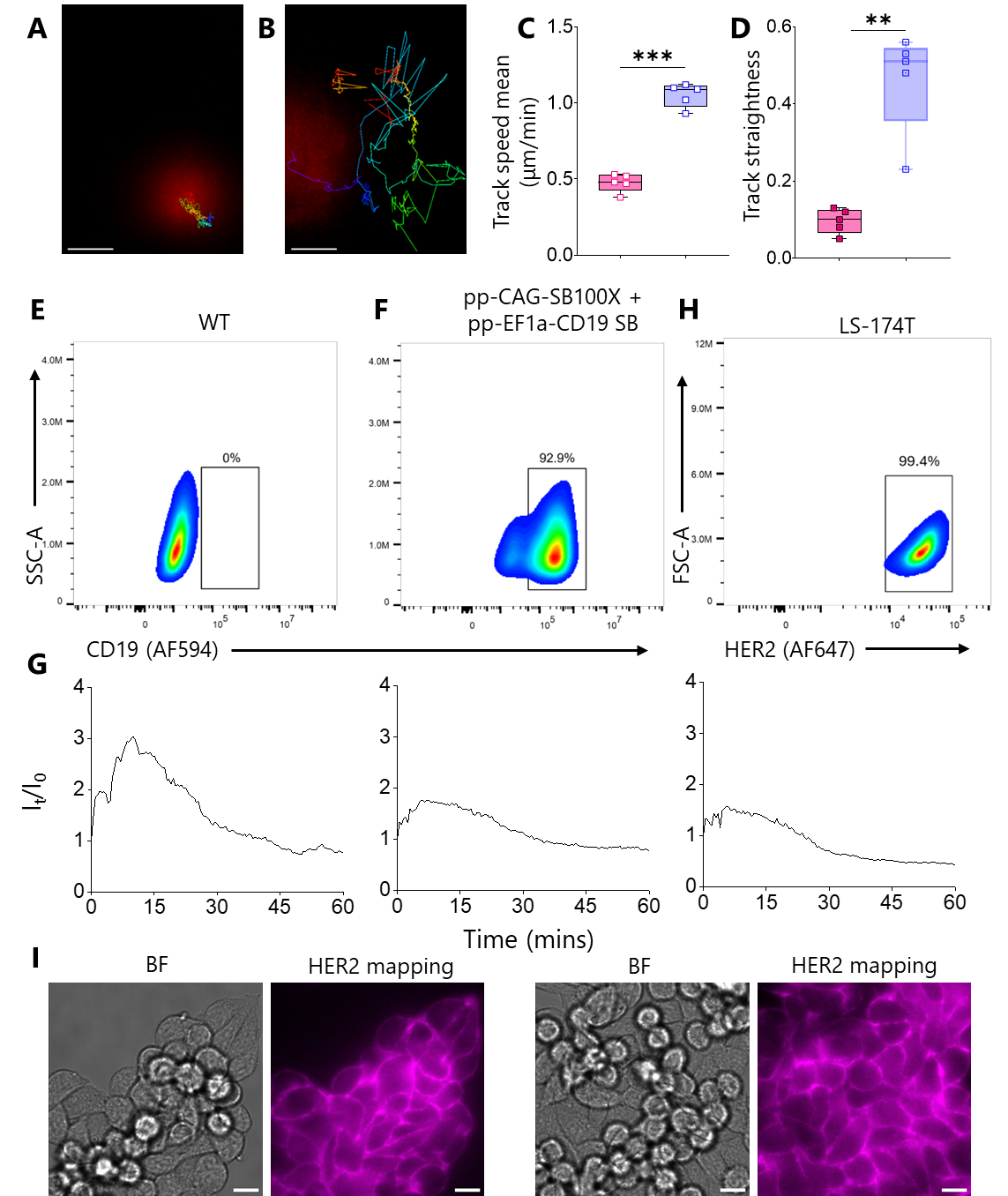

Fig. S3: (A-D) Representative tracks of CAR Jurkat T-cells on (A) early and (B) late seeded LS-174T monolayer. Scale bars are 4 μm. (C) Track speed mean (μm/min) and (D) track straightness of five individual T-cells on early (pink) and late (blue) seeded monolayers. Error bars depict mean ± SD. Means were compared using two-sided Student’s t-test. ***P = 0.0004, **P = 0.0018.

(E-F) Representative flow cytometry data of human CD19 expression in the LS-174T cell line. (E) CD19 expression in WT LS-174T cells (control). (F) 1 million LS-174T cells were electroporated with 250 ng of pp-CAG-SB100X and 1 µg of pp-EF1a-CD19 SB. 24-hours post nucleofection, cells were treated with 1 µg/mL puromycin. Following 1 week of puromycin selection, cells were stained for CD19 and analyzed by flow cytometry.

(G) Representative calcium flux traces of anti-CD19 Jurkat CAR T-cells interacting with late seeded CD19^+^ LS-174T monolayers.

(H) Representative flow cytometry data of HER2 expression in LS-174T.

(I) HER2 mapping on early (left) and late (right) seeded LS-174T. Scale bars are 10 μm.

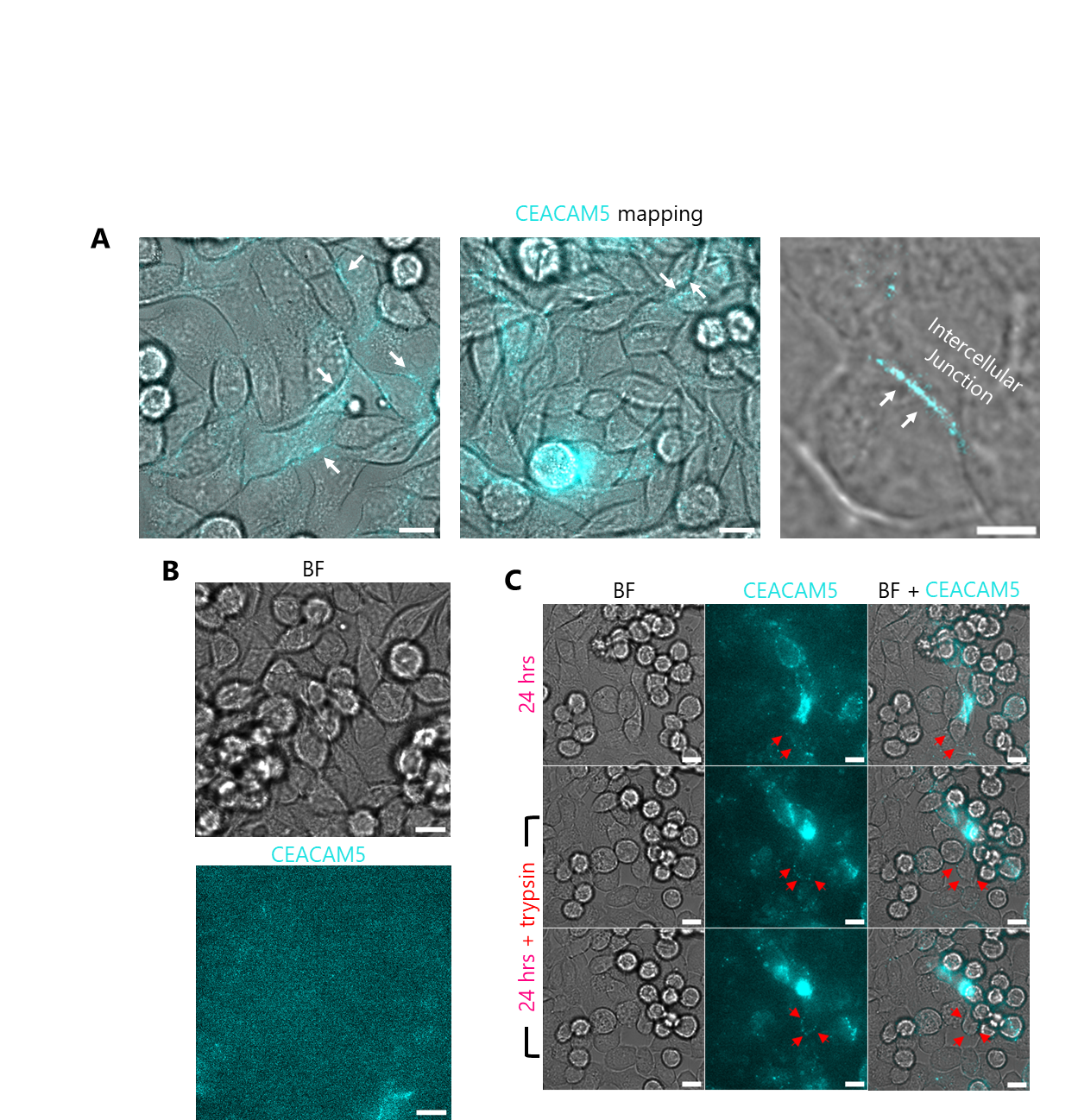

Fig. S4: (A) More images showing CEACAM5 is located at the intercellular junctions (white arrows) of early seeded LS-174T cells. Scale bars are 10 μm (left and middle images) and 5 μm (right image).

(B) CEACAM5 mapping on late seeded LS-174T monolayer. Scale bars are 10 μm.

(C) CEACAM5 mapping before (top row) and after (middle and bottom rows) trypsin treatment on early-seeded LS-174T monolayers. Red arrows show CEACAM5 staining on a single cell before and after the treatment. Scale bars are 10 μm.

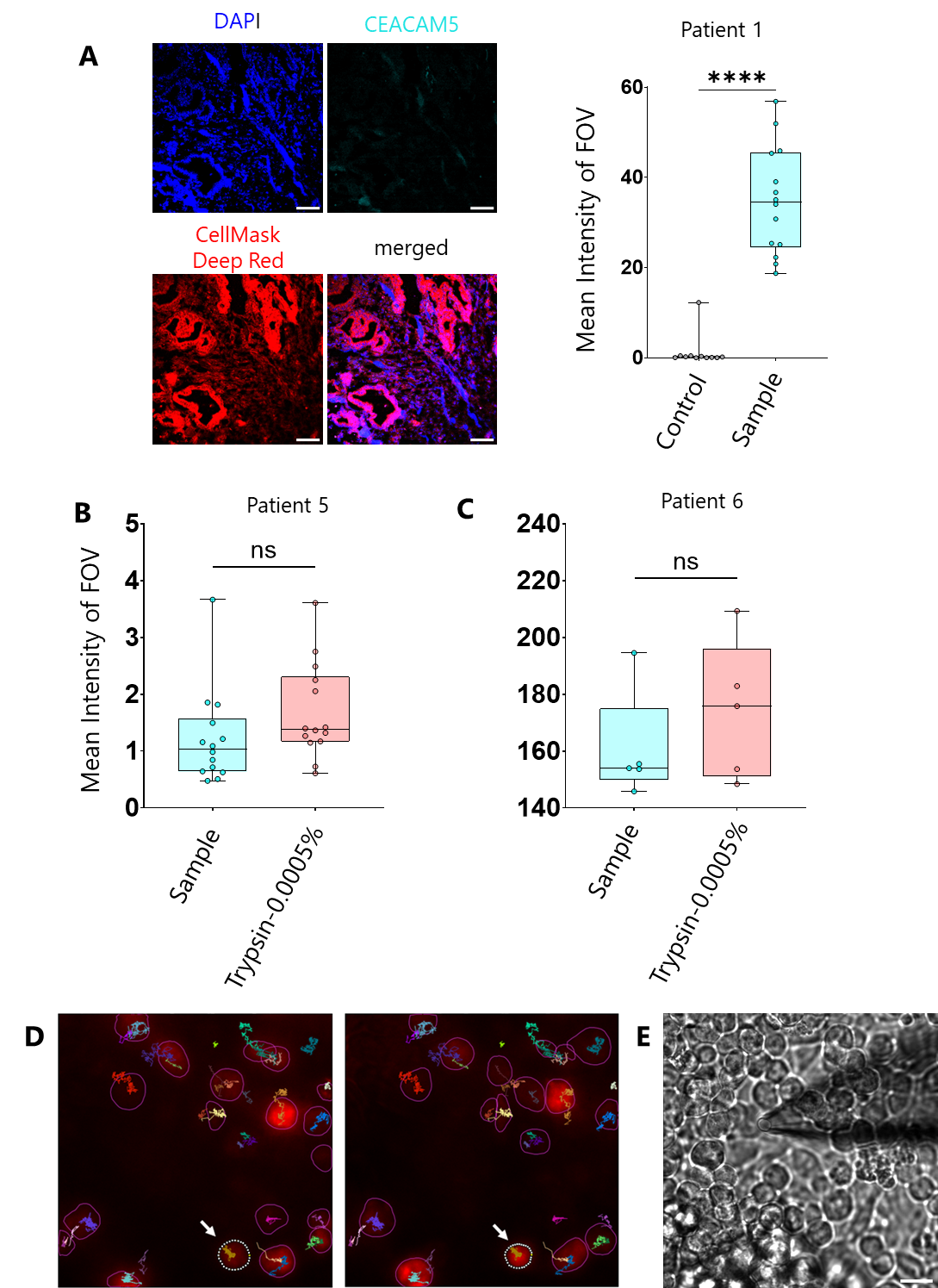

Fig. S5: (A) Representative control DAPI (blue), CEACAM5 (cyan), CellMask Deep Red (red) and merged images of patient 1 tissue stained with secondary antibody only. Scale bars are 50 μm. Boxplot shows the comparison of mean CEACAM5 intensity of patient 1 in control vs. sample.

(B & C) Comparison of mean CEACAM5 intensity between sample and trypsin (0.0005 %) treated conditions for patient 5 and 6 tissues. Medians in (A-C) were compared using two-sided Student’s t-test. ****P < 0.0001, ns not significant.

(D) Analysis steps of epifluorescence calcium imaging time-series movies. T-cells were detected using StarDist based on their cell shape. Extracted Mean fluorescence intensity (left panel: 14810, right panel: 25550) of two different frames are shown for representative cell (white arrow). All the frames were analyzed in similar way.

(E) Brightfield image of micropipette positioned above LS-174T cell monolayer prior to trypsin delivery. Tip diameter = 4-5 µm. Scale bar is 10 µm.

**Description of supplemental movies**

**Movie S1**

*Jurkat CAR T cell/SLB2 interaction*

Representative three-colour TIRF movie of CEACAM5 targeting Jurkat CAR T-cells interacting with CEACAM5^+^ and CEACAM5^-^ SLB2s. In each condition, three channels show CD45/CD43 exclusion on SLB2, T-cell footprint and calcium flux. Scale bar is 5 μm and the movie duration is 10 mins.

**Movie S2**

*Jurkat CAR T cell/LS-174T interaction*

Representative brightfield and epi-fluorescence calcium imaging movie of Jurkat CAR T-cells interacting with CEACAM5^+^ LS-174T cells. LS-174T cells were seeded for 24 hrs (early seeded, pink) and 48 hrs (longer seeded, blue) within incubator before the experiments. Red arrows indicate triggered Jurkat CAR T-cells on early seeded LS-174T monolayer. Scale bar is 20 μm and the movie duration is 60 mins.

**Movie S3**

*Jurkat CAR T-cell/LS-1034 interaction*

Representative brightfield and epi-fluorescence calcium imaging movie of Jurkat CAR T-cells interacting with early seeded CEACAM5^+^ LS-1034 cells. Scale bar is 20 μm and the movie duration is 60 mins.

**Movie S4 & S5**

*Jurkat CAR T-cell/CRC cell interaction*

Representative epi-fluorescence calcium imaging movie of Jurkat CAR T-cells interacting with early and longer seeded LS-174T (pink, blue) and early seeded RKO monolayer (olive). Colour coded calcium traces are shown in the movie. Scale bar is 5 μm and the original movie duration is 60 mins.

**Movie S6**

*Primary CAR T-cell/CRC cell interaction*

Representative brightfield movie of primary CAR T-cells interacting with early and longer seeded LS-174T and early seeded RKO monolayers. Red arrows indicate killing of LS-174T cells by primary CAR T-cells. Scale bar is 5 μm and the original movie duration is 60 mins.

**Movie S7 & S8**

*Primary CAR T-cell/CRC cell interaction*

Representative epi-fluorescence calcium imaging movie of primary CAR T-cells interacting with early and longer seeded LS-174T and early seeded RKO monolayers. Calcium traces are colour coded in the movies. Scale bar is 5 μm and the original movie duration is 60 mins.

**Movie S9**

*Tracking analysis of Jurkat CAR T-cells*

Imaris 10.0.1 (surface creation tool) was used for tracking the Jurkat CAR T-cells while interacting with early and longer seeded LS-174T monolayers. Surface is denoted by blue border and the dynamics of tracks are shown by the dragon tail representation (blue line). Scale bar is 10 μm and the movie duration is 60 mins.

**Movie S10**

*Granules release by primary CAR T-cells*

Representative epi-fluorescence imaging and brightfield movie of primary CAR T-cells interacting with early seeded LS-174T monolayer. Granules release at the immunological synapse (IS) is shown by white arrows (left movie). Scale bar is 5 μm and the original movie duration is 22 mins.

**Movie S11**

*Tracking analysis of released granules*

Imaris 10.0.1 was used for tracking analysis of released granules shown in the supplemental Movie S11. Surface is denoted by a light-yellow border and the dynamics of tracks are shown by the dragon tail representation (yellow line). Full tracks of the granules are shown in Fig. 3C (white arrows). Track speed mean of the granules was determined from this analysis and reported in Fig. 3D. Scale bar is 5 μm and the original movie duration is 22 mins.

**Movie S12**

*Trypsin treatment on LS-174T cells’ monolayer using a micropipette*

Representative brightfield movie shows local trypsin treatment on longer seeded LS-174T monolayer using a glass micropipette to dissociate the cells. Scale bar is 5 μm and time of treatment varies as required.

**Movie S13 & S14**

*Calcium imaging comparison of CAR T-cells before and after trypsin treatment*

Representative epifluorescence calcium imaging movie of Jurkat CAR (Movie S15) and primary CAR (Movie S16) T-cells interacting with longer seeded LS-174T monolayer before (blue) and after (orange) trypsin treatment. Colour coded calcium traces are shown in the movies. Scale bar is 5 μm and the movie duration is 60 mins.

**Movie S15**

*Increased killing of LS-174T cells after trypsin treatment*

Representative movie shows increased killing of LS-174T cells by primary CAR T-cells on longer seeded trypsin treated monolayer. The brightfield movie confirms the killing events (red arrows). Scale bar is 5 μm and the movie duration is 60 mins.

| Objective lens used | Movie no. |
| --- | --- |
| 20X | S2 & S3 |
| 60X | S4-S8, S10, S12-S15 |
| 100X | S1 |

Table 1: Objective lens used in different movies.
